# Triblock copolymer micelle model of spherical paraspeckles

**DOI:** 10.1101/2020.11.01.364190

**Authors:** Tetsuya Yamamoto, Tomohiro Yamazaki, Tetsuro Hirose

**Affiliations:** Institute for Chemical Reaction Design and Discovery, Hokkaido University, Kita 21, Nishi 10, Kita-ku, Sapporo, Hokkaido, Japan; PRESTO, Japan Science and Technology Agency (JST), 4-1-8 Honcho, Kawaguchi, Saitama, 332-0012, Japan; Graduate School of Frontier Biosciences, Osaka University, Suita 565-0871, Japan

## Abstract

Paraspeckles are nuclear bodies composed of architectural RNA (arcRNA) and RNA-binding proteins. In the wild type, the blocks at the two terminal regions of arcRNAs compose the shell of paraspeckles and the middle region between the two terminal blocks composes the core, analogous to micelles of ABC triblock copolymers. We here use an extension of the theory of polymer micelles to predict the structure and size of paraspeckles as one decreases the length of one of the terminal blocks of arcRNA by CRISPR/Cas9, assuming that paraspeckles are spherical. Our theory predicts that when the length of the edited terminal blocks is larger than a critical value, paraspeckles show discontinuous transitions between the structure in which all the edited terminal blocks are localized in the shell and the structure in which all the edited terminal blocks are localized in the core at a threshold value of the transcription rate of arcRNA. In contrast, when the length of the edited terminal blocks is smaller than the critical value, the population of edited terminal blocks in the shell decreases continuously as one increases the transcription rate of arcRNA. The size of paraspeckles increases as one decreases the length of the edited terminal blocks. These predictions are consistent with our experiments.

## INTRODUCTION

Cell nucleus is not a uniform solution of DNA and proteins: chromatin is distributed with non-uniform local concentration and there are a number of nuclear bodies (1, 2, 3, 4, 5, 6). Nuclear bodies are typically composed of RNA and RNA-binding proteins. Growing number of evidences suggest that nuclear bodies are assembled by phase separation due to the multi-valent interactions between the prion-like domains (or the intrinsically disordered regions) of proteins bound to RNA (1, 2, 3, 4, 5, 6). Direct RNA-RNA interactions due to base pairing may also be involved in the phase separation (6). Paraspeckles are nuclear bodies scaffolded by NEAT1 _2, which are long non-coding architectural RNAs (arcRNAs (1)) (7, 8, 9). These nuclear bodies function as molecular sponges that sequester some types of RNAs and proteins (2). Paraspeckles are often observed at the proximity to the transcription site of NEAT1 _2 and are disassembled when the transcription of the arcRNA is suppressed (7, 8, 10). The number of paraspeckles increase when transcription of NEAT1 _2 is upregulated (8, 13). These results imply that paraspeckles are assembled due to the production of NEAT1 _2 by transcription. RNA-binding proteins may be bound to nascent NEAT1 _2 transcripts, which are under production, and the interactions between these nascent NEAT1 _2 molecules may facilitate the nucleation and growth of paraspeckles (11, 12).

We have used an extension of the Flory-Huggins theory (which is the standard theory of phase separation of polymers in a solution (14)) to predict the phase separation due to the constant production of arcRNAs (15). This theory predicts that the condensates assembled by this mechanism are disordered liquids of arcRNAs associated by RNA-binding proteins. However, paraspeckles are not disordered liquid condensates of NEAT1 _2, but form the core-shell structure (16, 17): In wild type, the two terminal regions of NEAT1 _2 consist of the shell and the middle region consists of the core. This structure is analogous to micelles of ABC block copolymers in selective solvent, where the A and C blocks are hydrophilic and the B block is hydrophobic (18). The core-shell structure results from the fact that NEAT1 _2 transcripts are associated by proteins, such as NONO and FUS, bound in the middle region of NEAT1 _2 (19). The ordered structure and the transcription driven formation of paraspeckles are in contrast to condensates assembled by the classical liquid-liquid phase separation in the equilibrium.

In our recent experiment, we have produced constructs, with which part of the terminal regions of NEAT1 _2 transcripts was deleted by CRISPR/Cas9, and paraspeckles assembled in such constructs were observed by super-resolution optical microscope and electron microscope (19, 20). The edited terminal regions are distributed to the paraspeckle core as well as to the shell. Our experiments have shown that the distribution of the terminal regions and the size of paraspeckles depend on the length of the terminal regions (19, 20). Motivated by this experiment, we here model a paraspeckle as a micelle of ABC triblock copolymers and theoretically predict the distribution of terminal blocks and the size of paraspeckles as the length of one of the two terminal blocks (A blocks) decreases. This paper treats only cases in which paraspeckles are spherical.

Our theory predicts that when A blocks are longer than a critical value, either the ‘all-shell structure’ with which all the A blocks are in the shell or the ‘all-core’ structure with which all the A blocks are in the core is stable. There is a discontinuous transition between the two structures at a threshold value of the production rate of NEAT1 _2. It is because when the length of A blocks is large, the free energy due to the excluded volume interactions between A blocks in the shell and the free energy due to the excluded volume interactions between A and B blocks in the core dominate the mixing free energy, analogous to the Flory-Huggins theory. In contrast, when the length of A blocks is smaller than the critical value, the fraction of A blocks in the shell decreases continuously with increasing the production rate of NEAT1 _2 (mixed structure). The size of paraspeckles increases as the length of A blocks decreases (while the production rate of NEAT1 _2 is fixed). It is because the number of NEAT1 _2 in a paraspeckle is limited by the excluded volume interactions between A blocks in the shell and the free energy due to the latter interactions decreases as the length of A blocks decreases. These predictions are consistent with the results of our experiments (19, 20).

## MATERIALS AND METHODS

### Free energy

We use an extension of the theory of block copolymer micelles (21, 22, 23) to treat paraspeckles. For simplicity, we here treat spherical paraspeckles, although cylindrical paraspekcles with aspect ratio 1 − 2 are often observed for cases in which one terminal block is deleted significantly. NEAT1_2 transcripts that compose a paraspeckle is modeled as ABC triblock copolymer, see fig. 1**a**. A, B, and C blocks are composed of *N*_A_, *N*_B_, and *N*_C_ Kuhn segments. The length of each Kuhn segment is *b*. B blocks compose the core and C blocks compose the shell. The fraction *α* of A blocks is distributed in the shell and the others are distributed in the core. For simplicity, we take into account RNA-binding proteins only implicitly. This assumption is effective for the case in which the binding affinity of RNA-binding proteins to NEAT1_2 is relatively large (15).

**Figure 1.**
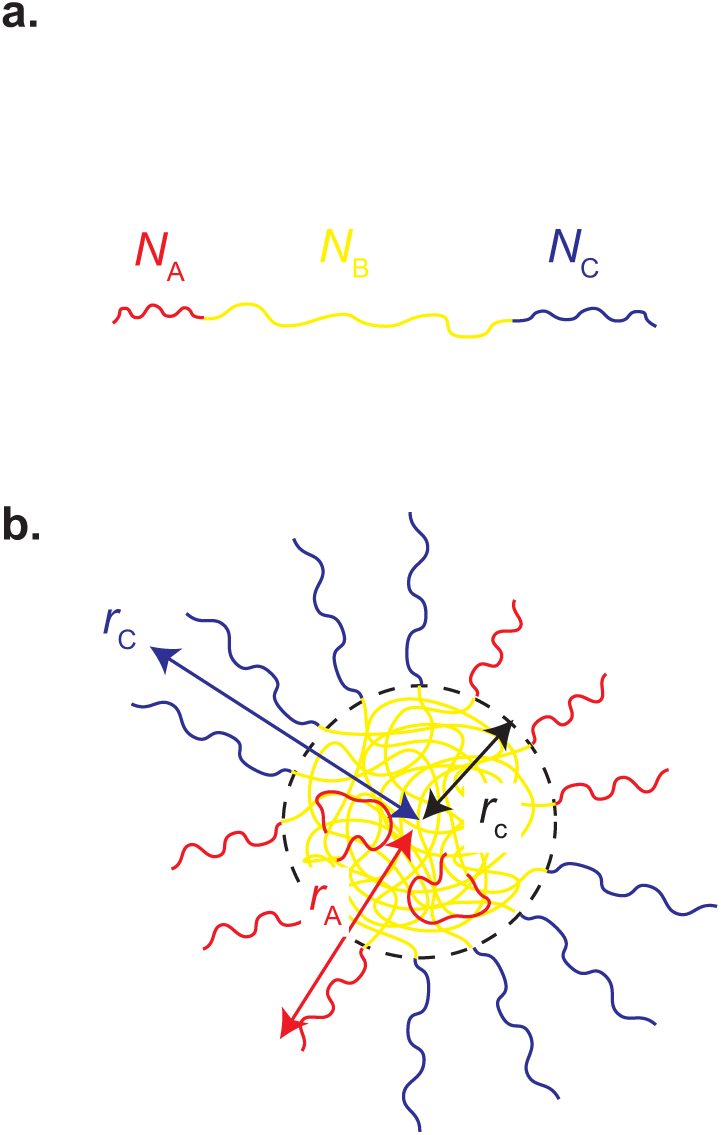
Spherical paraspeckles are modeled as micelles of ABC block copolymers. A, B, and C blocks are composed of *N*_A_, *N*_B_, and *N*_C_ segments (**a**). Each paraspeckle is composed of *n* copolymers. B blocks of copolymers are packed in the core of the paraspeckle and C blocks are localized in the shell (**b**). A fraction *α* of A blocks is localized at the shell and the other fraction is in the core. A and C blocks are located in distinct domains in the shell. The size of the paraspeckle is characterized by the radius *r*_c_ of the core, the distance *r*_A_ between the top of an A domain and the center of the paraspeckle, and the distance *r*_C_ between the top of a C domain and the center of the paraspeckle.

The free energy of a paraspeckle composed of *n* triblock copolymers has the form

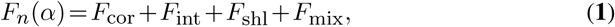

where *F*_cor_ is the free energy of the core, *F*_int_ is the interfacial free energy, *F*_shl_ is the free energy of the shell, and *F*_mix_ is the mixing free energy.

The free energy of the core has the form

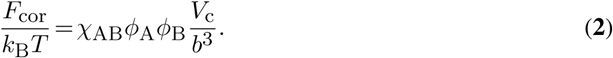

The interaction parameter *χ*_AB_ accounts for the excluded volume interactions between A blocks and B blocks. *V*_c_ is the volume of the core. *ϕ*_A_ (= *b*^3^(1 *- α*)*N*_A_*n/V*_c_) and *ϕ*_B_ (= *b*^3^*N*_B_*n/V*_c_) are the volume fraction of A and B blocks in the core, respectively. The core is occupied by A and B blocks, *ϕ*_A_ +*ϕ*_B_ = 1. This leads to the form of the volume

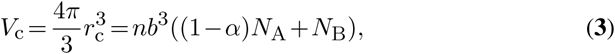

where *r*_c_ is the radius of the core. We assume that A blocks are distributed randomly in the core, motivated by experiments that suggest that the 3′ terminal regions of Δ3′-NEAT1_2 are distributed randomly in the core of paraspeckles. We also neglected the free energy contribution due to the conformational entropy of B blocks because this contribution is relatively small, see also the Supplementary File.

The interfacial free energy has the form

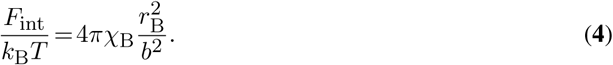

The interaction parameter *χ*_B_ accounts for the interactions between B blocks. Eq. (**4**) is the free energy cost due to the fact that B blocks at the interface cannot interact with other B blocks.

The free energy of the shell has the form

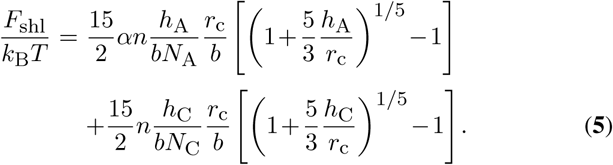

We used the mean field approximation that neglects the correlation due to the excluded volume interactions between segments in each A block and those between segments of each C block to derive eq. (**5**). The derivation of eq. (**5**) and the results by using the scaling theory are shown in the Supplementary File. We treat the blocks in the shell as a polymer brush as the usual treatment of polymer micelles (21, 22, 23). The 3′ and 5′ terminal regions are segregated in separate domains in the shell of wild type paraspeckles (17). Motivated by this experimental result, we do not take into account the excluded volume interactions between A and C blocks in the shell. Eq. (**5**) takes into account the free energy contributions due to the conformational entropy of A and C blocks, the free energy due to the excluded volume interactions between A blocks, and the free energy due to the excluded volume interactions between C blocks. *h*_A_ and *h*_C_ are the heights of A and C blocks in the limit of planer brush, *r*_c_ →∞, and have the forms (24, 25)

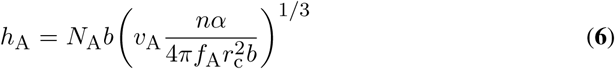

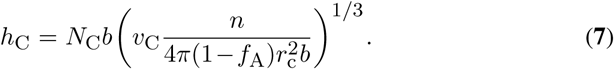

Eq. (**5**) returns to the free energy of planer brushes for *h*_A_ ≪*r*_c_ and *h*_C_≪ *r*_c_. The excluded volume *v*_A_ accounts for the excluded volume interactions between A blocks and the excluded volume *v*_C_ accounts for the excluded volume interactions between C blocks. We assume that the interactions between A blocks and the interactions between C blocks are repulsive, following the usual treatment of diblock copolymer micelles (21, 22, 23). We discuss this assumption in the Discussion section. The fraction *f*_A_ of the interface occupied by A blocks has the form

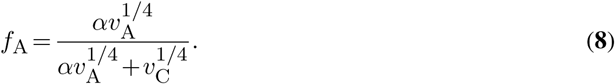

Eq. (**8**) is derived by using the fact that the osmotic pressure in the domains of A blocks is equal to the osmotic pressure in the domains of C blocks, see the Supplementary Materials.

The mixing free energy has the form

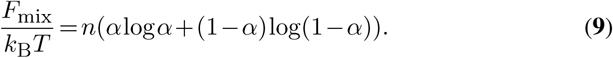

The free energy *F*_*n*_(*α*) is a function of the number *n* of triblock copolymers comprising the paraspeckle and the fraction *α* of A blocks in the shell. The fraction *α* is determined by the minimization of the free energy. This corresponds to cases in which the time scale of the redistribution of A blocks is smaller than the time scale of the production of new copolymers.

### Association and dissociation dynamics of arcRNA transcripts

Paraspeckles are assembled by the production of NEAT1 _2. The attractive interactions between nascent NEAT1 _2 transcripts, where their production is still in progress, probably facilitate the formation and growth of paraspeckles. We thus treat the dynamics with which NEAT1 _2 transcripts are added to the paraspeckle one by one with a constant rate. This is an extension of the classical theory of the dynamics of micellization (26).

We treat the probability *q*_*n*_(*t*) that the paraspeckle growing at the transcription site of NEAT1 _2 is composed of *n* triblock copolymers at time *t* after a transcription burst starts. The time evolution of the probability *q*_*n*_(*t*) has the form

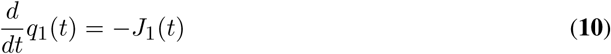

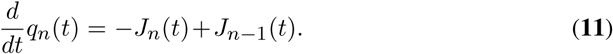

The flux *J*_*n*_(*t*) has the form

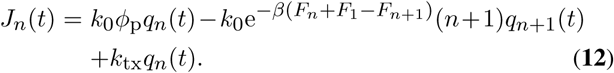

The first term of eq. (**12**) is the rate with which a triblock copolymer in the solution is spontaneously associated to the paraspeckle. The second term is the rate with which a triblock copolymer is spontaneously dissociated from the paraspeckle. The third term is the rate with which a triblock copolymer is added to the paraspeckle by transcription.

*k*_0_ is the inverse of the time scale of the thermal motion of triblock copolymers. *ϕ*_p_ is the volume fraction of triblock copolymers that are not associated to paraspeckles. *F*_*n*_ is the free energy which is already minimized with respect to the fraction *α*, see eq. (**1**). *k*_tx_ is the rate with which a nascent transcript is added to the paraspeckle. We note that eqs.(**10**) and (**11**) treat the dynamics of the probability that the paraspeckle growing at the transcription site of NEAT1 _2 is composed of *n* triblock copolymers. It is in contrast to many theories of micellation dynamics that describes the dynamics of the volume fraction of micelles composed of *n* polymers in a solution. When the transcription is stopped, *k*_tx_ → 0, the probability 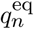 in the equilibrium has the form

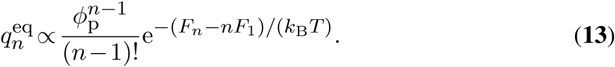

The forms of the first and second terms of eq. (**12**) thus ensures the detailed balance.

In the steady state, *dq*_*n*_(*t*)*/dt* = 0, the solution of eq. (**10**) and (**11**) has the form

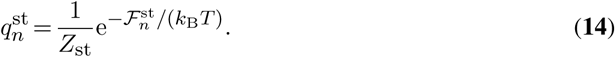

The effective free energy 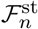 has the form

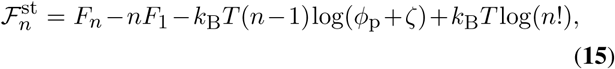

where *ζ* (= *k*_tx_*/k*_0_) is the ratio of the association rates. *Z*_st_ is the effective partition function

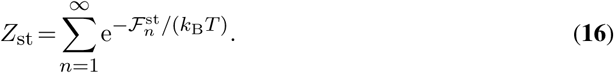

The number of triblock polymers composing a paraspeckle with the maximum probability in the steady state is derived by minimizing the effective free energy 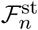 with respect to *n*.

In a more precise treatment, we need the time evolution equation of the volume fraction *ϕ*_p_ of unassociated triblock copolymers in the solution to close the theory. For simplicity, we here treat the case in which the volume fraction *ϕ*_p_ is small and the spontaneous association process is dominated by the incorporation of nascent transcripts by transcription. In this case, paraspeckles do not grow by coalescence even in the long time scale, in contrast to the micellization in a solution of amphiphiles (27, 28),

## RESULTS

Our theory predicts the fraction *α* of A blocks in the shell and the number *n* of triblock copolymers in a paraspeckle in the steady state as a function of the logarithm of the production rate log(*k*_tx_*/k*_0_) of copolymers (log is the natural logarithm), the number of segments of blocks (*N*_A_, *N*_B_, and *N*_C_), and interaction parameters (*v*_A_*/b*^3^, *v*_C_*/b*^3^, *χ*_B_, and *χ*_AB_). When the number of segments *N*_A_ of A blocks is smaller than a critical value *N*_Ac_, the fraction *α* decreases continuously with increasing the production rate of copolymers, see the cyan and light green lines in fig. 2. When the number of segments *N*_A_ of A blocks is larger than the critical value *N*_Ac_, paraspeckles show the ‘all-shell’ state, in which all the A blocks are in the shell, *α* ≃ 1, for small copolymer production rate and the ‘all-core’ state, in which all the A blocks are in the core, *α* ≃ 0, for large copolymer production rate, see the organge and magenta lines in fig. 2. There is a discontinuous transition between the all-shell and all-core states at a threshold value of copolymer production rate.

**Figure 2.**
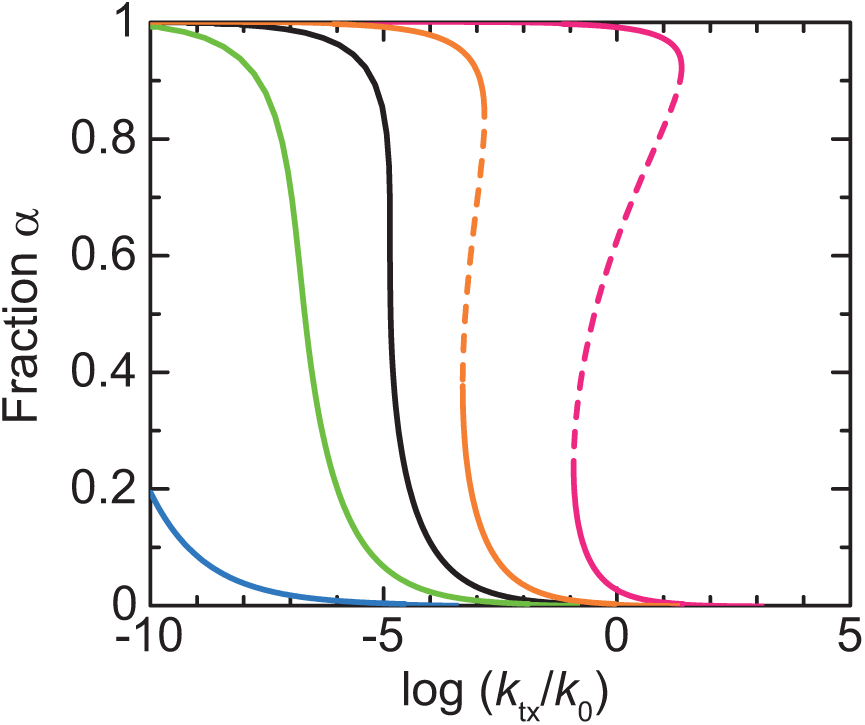
The fraction *α* of A blocks in the shell is shown as a function of the (natural) logarithm of the production rate log(*k*_tx_*/k*_0_) of NEAT1 _2. We used *N*_A_ = 5.0 (cyan), 10.0 (light green), 12.5228 (black), 15.0 (orange), and 20.0 (magenta) for the number of segments *N*_A_ of A blocks. The values of other parameters used for the calculations are *N*_B_ = 40.0, *N*_C_ = 20.0, *v*_A_*/b*^3^ = 1.0, *v*_C_*/b*^3^ = 1.0, *χ*_B_ = 0.5, and *χ*_AB_ = 1.0.

The fraction *α* of A blocks is determined mainly by the competition between the free energy due to the repulsive interactions between A blocks in the shell, the free energy due to the repulsive interactions between A and B blocks in the core, and the mixing free energy. The free energy of A blocks in the shell and core increases as the number of segments *N*_A_ of A blocks increases, whereas the mixing free energy is independent of the number of segments *N*_A_ of A blocks. When the number of segments *N*_A_ is larger than the critical value *N*_Ac_, the free energy of A blocks in the shell and core dominates the mixing free energy. In such situation, paraspeckles in either the all-shell state or all-core state are stable, see fig. 3. In contrast, when the number of segments *N*_A_ of A blocks is smaller than the critical value *N*_Ac_, the mixing free energy is significant and thus the mixed state, where A blocks are distributed between the shell and core, is stable, see fig. 3. This situation may be analogous to the Flory-Huggins theory that predicts that the translational entropy of polymers is suppressed due to their connectivity. The number *n* of triblock copolymers increases with increasing the production rate of copolymers. In the limit of large production rate of copolymers, all-core state becomes stable because the free energy due to the repulsive interactions between A and B blocks in the core increases as ∼*n* (see eq. (**2**)) while the free energy due to the excluded volume interactions between A blocks in the shell increases as ∼ *n*^11*/*9^ (see eq. (**5**)). The discontinuous transition between the all-shell state and the all-core state therefore results from the fact that the free energy is minimized by distributing A blocks in the shell for cases in which the number *n* of copolymers is small and by distributing A blocks in the core for cases in which the number *n* of copolymers is large. In the limit of small copolymer production rate, stable paraspeckles are not assembled, see the region delineated by the red line in fig. 3.

**Figure 3.**
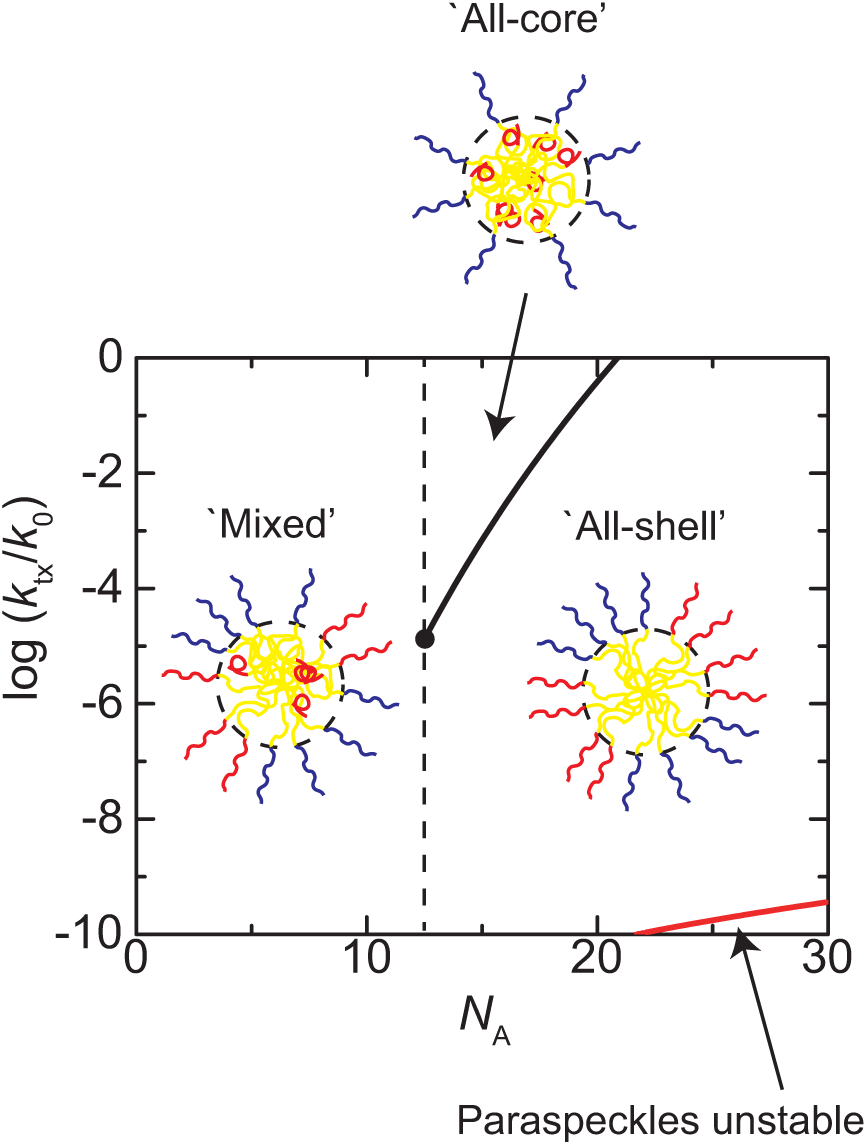
The phase diagram of paraspeckles is shown for the parameter space of the (natural) logarithm of production rate log(*k*_tx_*/k*_0_) of copolymers and the number of segments of A blocks *N*_A_. We used *N*_B_ = 40.0, *N*_C_ = 20.0, *v*_A_*/b*^3^ = 1.0, *v*_C_*/b*^3^ = 1.0, *χ*_B_ = 0.5, and *χ*_AB_ = 1.0. Paraspeckles are not stable for the copolymer production rate smaller than the red line.

When the production rate of copolymers is smaller than the critical value (log(*k*_tx_*/k*_0_) < − 4.8631 in fig. 3), the radius of paraspeckles (defined by the distance between the top of A or C domains and the center of the paraspeckle) increases with decreasing the number of segments *N*_A_ of A blocks, see fig 4. The average number *n* of block copolymers increases accordingly with decreasing the number of segments *N*_A_ blocks, see fig. 5. The number of copolymers in a paraspeckle is limited by the excluded volume interactions between A blocks (and the excluded volume interactions between C blocks) in the shell, analogous to micelles of diblock copolymers. The fraction *α* of A blocks in the shell decreases with decreaing the number of segments *N*_A_ of A blocks, see fig. 6. More copolymers can be therefore incorporated to paraspeckles as the number of segments *N*_A_ of A blocks decreases. Indeed, the number *n* of copolymers in a paraspeckle does not increase significantly with decreasing the number of segments *N*_A_ of A blocks for *N*_A_ > *N*_Ac_, where the fraction *α* is unity. The number *n* of copolymers in a paraspeckle increases steeply for *N*_A_ < *N*_Ac_, where the fraction *α* decreases steeply.

**Figure 4.**
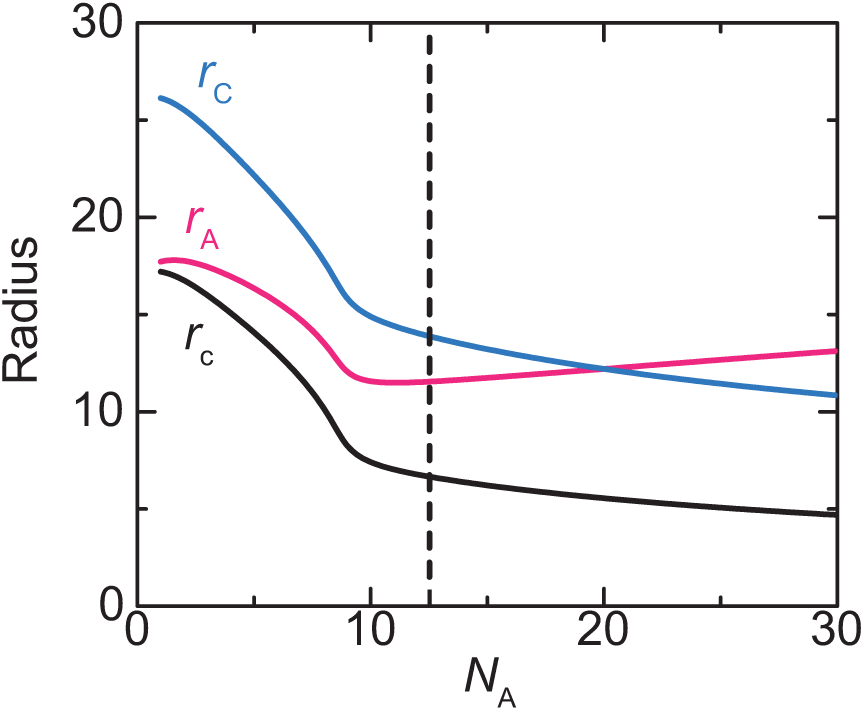
The radius *r*_c_ of the core of a paraspeckle (black), the distance *r*_A_ between the top of A domains and the center of the paraspeckle (magenta), and the distance *r*_C_ between the top of C domains and the center of the paraspeckle (cyan) are shown as functions of the number of segments *N*_A_ of A blocks. The values of parameters used for the calculations are log(*k*_tx_*/k*_0_)= −8.0, *N*_B_ = 40.0, *N*_C_ = 20.0, *v*_A_*/b*^3^ = 1.0, *v*_C_*/b*^3^ = 1.0, *χ*_B_ = 0.5, and *χ*_AB_ = 1.0. The critical number of segments of A blocks, *N*_Ac_, is indicated by the broken line.

**Figure 5.**
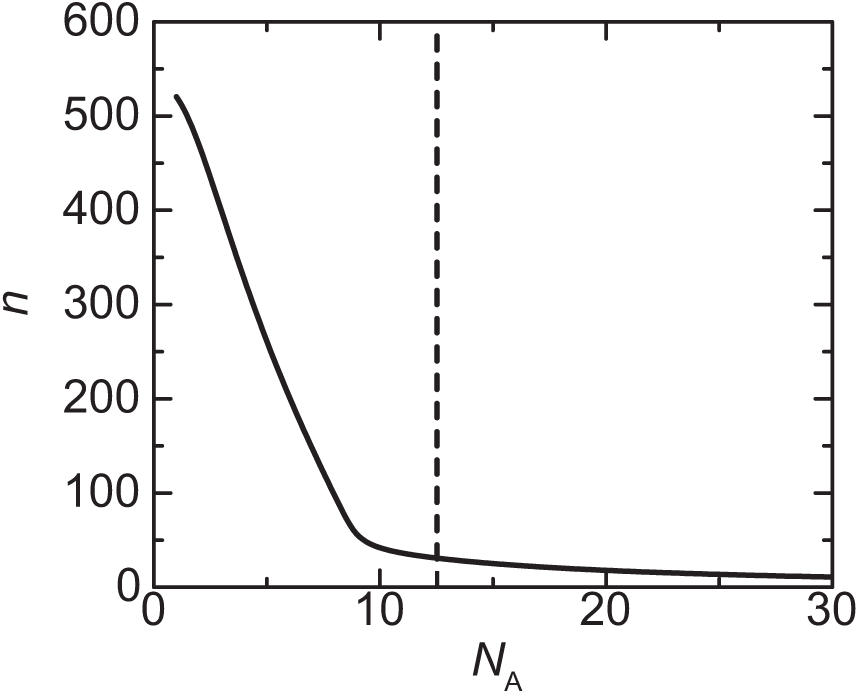
The number *n* of copolymers of a paraspeckle in the steady state is shown as a function of the number of segments *N*_A_ of A blocks. The values of parameters used for calculations are log(*k*_tx_*/k*_0_)= −8.0, *N*_B_ = 40.0, *N*_C_ = 20.0, *v*_A_*/b*^3^ = 1.0, *v*_C_*/b*^3^ = 1.0, *χ*_B_ = 0.5, and *χ*_AB_ = 1.0. The critical number of segments of A blocks, *N*_Ac_, is indicated by the broken line.

**Figure 6.**
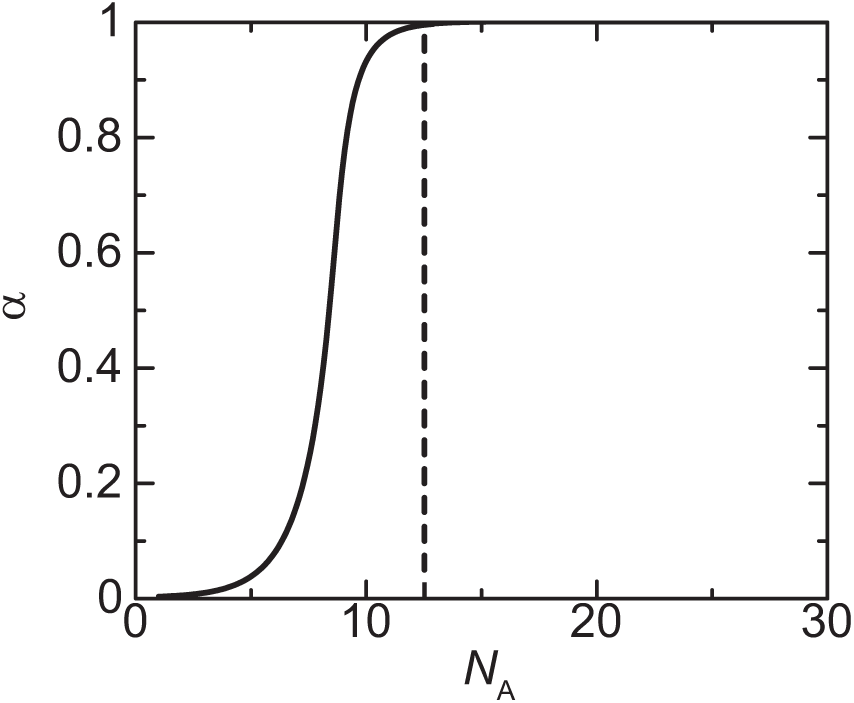
The fraction *α* of A blocks in the shell is shown as a function of the number of segments *N*_A_ of A blocks. The values of parameters used for calculations are log(*k*_tx_*/k*_0_)= −8.0, *N*_B_ = 40.0, *N*_C_ = 20.0, *v*_A_*/b*^3^ = 1.0, *v*_C_*/b*^3^ = 1.0, *χ*_B_ = 0.5, and *χ*_AB_ = 1.0. The critical number of segments of A blocks, *N*_Ac_, is indicated by the broken line.

## DICUSSION

We have used an extension of the theory of ABC triblock copolymer micelles to predict the structure of paraspeckles as a function of the production rate of copolymers and the number of segments *N*_A_ of A blocks. Decreasing the number of segments *N*_A_ of A blocks corresponds to deleting 3′ or 5′ terminal regions of NEAT1 _2. Our theory predicts that when the number of segments *N*_A_ of A blocks is larger than the critical value *N*_Ac_, either the all-shell state or all-core state is stable. It is because the free energy due to the excluded volume interactions between A blocks in the shell and the free energy due to the excluded volume interactions between A blocks and B blocks increases with increasing the number of segments *N*_A_ of A blocks, while the mixing free energy is independent of the number of segments *N*_A_ of A blocks. This concept is analogous to the Flory-Huggins theory. For relatively large values of interaction parameter *χ*_AB_, all-shell state is stable for small copolymer production rate and all-core state is stable for large copolymer production rate. Paraspeckles thus show a discontinuous transition at a threshold value of copolymer production rate. These results reflect the features of the free energy of NEAT1 _2 copolymers, eq. (**1**), rather than the production dynamics of copolymers. This prediction is thus also effective to micelles of ABC triblock copolymers, where A and C blocks are separated into different domains in the shell.

Our present model takes into account two features of paraspeckles: First, paraspeckles form the core-shell structure. We use an extension of the free energy of block copolymer micelles to take into account the core-shell structure of paraspeckles. This model assumes the core-shell structure and spherical shapes of paraspeckles. Taking into account the long-range interactions by using the Ohta-Kawasaki free energy (29, 30) and/or the free energy of density functional theory (31) in an extension of our previous theory of phase separation by the production dynamics of arcRNA transcripts (15) may be a complementary approach. Indeed, paraspeckles are cylindrical with aspect ratio of 1 - 3 and the aspect ratio depends on the length of 3’ and 5’ terminal blocks. Theories of polymer micelles predict that the conformational entropy of B blocks plays a significant role in the spherical-cylindrical transitions of micelles, albeit not in their size (21, 22, 23). Our theory can be extended to cylindrical paraspeckles by taking into account the conformational entropy of B blocks in the core and by analyzing the time evolution of the size of paraspeckles. Second, paraspeckles are produced by transcription dynamics. We use a simple model to take into account the process with which nascent NEAT1 _2 transcripts, under production, are incorporated one by one to the growing paraspeckle. Indeed, more direct experimental evidences of the latter process remain to be collected. Nevertheless, with this process, nascent NEAT1 _2 transcripts are incorporated to the growing paraspeckles without translational entropy cost and the relaxation time of paraspeckles (until a growing paraspeckle diffuses out from the transcription site) is enhanced because the paraspeckle is connected to the transcription site via the nascent transcripts and RNA polymerases. It is of interest to take into account the contributions of the nascent transcripts and RNA polymerases in an extension of our theory.

Our predictions result from the assumptions that 1) A and C blocks are localized in separate domains in the shell and 2) the interactions between A blocks and those between C blocks are repulsive. Indeed, super-resolution optical microscopes have elucidated that there are multiple A and C domains in the shell of each paraspeckle and that paraspeckles form by two-step process − NEAT1 _2 transcripts form bundles and these bundles associate to form paraspeckles (17). It is thus of interest to predict the pattern of A and C blocks in an extension of our theory. With our theory, the result that the size of paraspeckles increases with decreasing the length of A blocks is owing to the excluded volume interactions between blocks in the shell. Indeed, the coalescence of paraspeckles has not been observed even when paraspeckles of wild type NEAT1 _2 are close vicinity, implying that the repulsive interactions between A blocks and those between C blocks suppress the coalescence. Critical experimental tests on the repulsive interactions may greatly advance our understanding of the mechanism behind the assembly of paraspeckles.

Some predictions of our theory may be experimentally accessible: 1) As the length *N*_A_ of one of the two terminal regions of NEAT1 _2 decreases, the fraction of the terminal regions in the shell of paraspeckles decreases and the size of paraspeckles increases. 2) The fraction of the terminal regions in the shell of wild type paraspeckles does not change significantly with increasing the transcription rate of NEAT1 _2, unless the transcription rate becomes larger than the threshold value of the discontinuous transition. In contrast, the fraction of the terminal regions in the shell of paraspeckles of end-deleted NEAT1 _2 decreases continuously with increasing the transcription rate of NEAT1 _2. Here, the transcription rate refers to the production rate of NEAT1 _2 transcripts in one transcription burst. Increasing the frequency of transcription burst probably increases the number of paraspeckles. The frequency of transcription burst may affect the size of paraspeckles and the fraction of the terminal regions of NEAT1 _2 when it increases the concentration of NEAT1 _2 at the vicinity of transcription site of NEAT1 _2 significantly.

Our recent experiment reveals that 1) the 5^′^ terminal regions of Δ5′ constructs, where the 5′ terminal regions of NEAT1 _2 is deleted by ≈2 kbps, are distributed randomly between the shell and the core, 2) most of the 3′ terminal regions of Δ3′ constructs, where the 3′ terminal regions of NEAT1 _2 is deleted by 6 kbps, are localized in the core, 3) the sizes of paraspeckles of Δ5′ and Δ3′ construts are larger than the size of paraspeckles of wild type NEAT1 _2 (20). The results of our theory are therefore in consistent with experiments. We therefore conclude that paraspeckles are modeled as ABC triblock copolymer micelles, at least, to the first approximation. Our experiments suggest that RNA-binding proteins may be localized in specific RNA regions, although our theory treats that each block is uniform. It is of interest to take into accout the code imprinted in the sequence of bases of NEAT1 _2 in an extension of our theory.

## ACKNOWLEDGEMENTS

This work was supported by JST, PRESTO Grant Number JPMJPR18KA and JSPS Kakenhi Grant Number 19H05259 and 18K03558 (PI: T. Yamamoto) and 19H05250 and 19K06479 (PI: T. Yamazaki). Tetsuya Yamamoto is grateful to the fruitful discussion with Takashi Uneyama (Nagoya University).

## REFERENCES

1. Chujo, T., Yamazaki, T., and Hirose, T. (2016) Architectural RNAs (arcRNAs): A class of long noncoding RNAs that function as the scaffold of nuclear bodies. Biochim. Biophys. Acta, 1859, 139–146.

2. Nakagawa, S., Yamazaki, T., and Hirose, T. (2018) Molecular dissection of nuclear paraspeckles: towards understanding the emerging world of the RNP milieu. Open Biol., 8, 180150.

3. Palikyaras, S. and Papantonis, A. (2019) Modes of phase separation affecting chromatin regulation. Open Biol., 9, 190167.

4. Banani, S.F., Lee, H.O., Hyman, A.A., and Rosen, M.K. (2017) Biomolecular condensates: organizers of cellular biochemistry. Nat. Rev. Mol Cell Biol., 18, 285–298.

5. Berry, J., Weber, S.C., Vaidya, N., Haataja, M., and Brangwynne, C.P. (2015) RNA transcription modulates phase separation-driven nuclear body assembly. Proc. Nat. Acad. Sci. USA, 112, E5237–E5245.

6. Van Treeck, B. and Parker, R. (2018) Emerging Roles for Intermolecular RNA-RNA Interactions in RNP Assemblies.. Cell, 174, 791–802.

7. Sasaki, Y.T.F., Ideue, T., Sano, M., Mituyama, T., and Hirose, T. (2009) MEN*ϵ/β* noncoding RNAs a re essential for structural integrity of nuclear paraspeckles. Proc. Nat. Acad. Sci. USA, 106, 2525–2530.

8. Clemson, C.M., Hutchinson, J.N., Sara, S.A., Ensminger, A.W., Fox, A.H.,Chess, A., and Lawrence, J.B. (2009) An Architectural Role for a Nuclear Noncoding RNA: NEAT1 RNA Is Essential for the Structure of Paraspeckles. Cell, 33, 717–726.

9. Sunwoo, H., Dinger, M.E., Wilusz, J.E., Amaral, P.P., Mattick, J.S., and Spector, D.L. (2008) MEN *ϵ/β* nuclear-retained non-coding RNAs are up-regulated upon muscle differentiation and are essential components of paraspeckles. Genome Res., 19, 347–359.

10. Mao, Y.S., Sunwoo, H., Zhang, B., and Spector, D.L. (2011) Direct visualization of the co-transcriptional assembly of a nuclear body by noncoding RNAs. Nat. Cell Biol., 13, 95–101.

11. Chujo, T. and Hirose, T. (2017) Nuclear Bodies Built on Architectural Long Noncoding RNAs: Unifying Principles of Their Construction and Function. Mol. Cell, 40, 889–896.

12. Yamazaki, T., Nakagawa, S., and Hirose, T. (2020) Architectural RNAs for Membraneless Nuclear Body Formation. Cold Spring Harb. Symp. Quant. Biol., 10.1101/sqb.2019.84.039404.

13. Hirose, T., Virnicchi, G., Tanigawa, A., Naganuma, T., Li, R., Kumura, H., Yokoi, T., Nakagawa, S., Bénard, M., Fox, A.H., and Pierron, G., (2014) NEAT1 long noncoding RNA regulates transcription via protein sequestration within subnuclear bodies. Mol. Biol. Cell, 25, 169–183.

14. Doi, M. (1996) Introduction to Polymer Physics. Oxford Univ. Press, Oxford, New York, US.

15. Yamamoto, T., Yamazaki, T., Hirose, T. Phase separation driven by production of architectural RNA transcripts. Soft Matter, 10.1039/C9SM02458A.

16. Souquere, S., Beauclair, G., Harper, F., Fox, A., and Pierron, G. Highly Ordered Spatial Organization of the Structural Long Noncoding Neat1 RNAs Within Paraspeckle Nuclear Bodies. (2010) Mol. Biol. Cell, 21, 4020–4027.

17. West, J.A., Mito, M., Kurosaka, S., Takumi, T., Tanegashima, C., Chujo, T., Yanaka, K., Kingston, R.E., Hirose, T., Bond, C., Fox. A., and Nakagawa, S. (2016) Structural, super-resolution microscopy analysis of paraspeckle nuclear body organization. J. Cell Biol., 214, 817–830.

18. Monzen, M., Kawakatsu, T., Doi, M., Hasegawa, R. (2000) Micelle formation in triblock copolymer solutions Comput. Theor. Polym. S., 10, 275–280.

19. Yamazaki, T., Souquere, S., Chujo, T., Kobelke, S., Chong, Y.S., Fox, A.H., Bond, C.S., Nakagawa, S., Pierron, G., and Hirose, T. (2018) Functional Domains of NEAT1 Architectural IncRNA Induce Paraspeckle Assembly through Phase Separation. Mol. Cell, 70, 1038–1053.

20. Yamazaki, T., et al. (2020) Title BioArxiv, doi

21. Halperin, A. and Alexander, S. (1989) Polymeric Micelles: Their Relaxation Kinetics. Macromolecules, 22, 2403–2412.

22. Semenov, A.N., Nyrkova, I.A., and Khokhlov, A.R. (1995) Polymers with Strongly Interacting Groups: Theory for Nonspherical Multiplets. Macromolecules, 28, 7491–7500.

23. Zhulina, E.B., Adam, M., LaRue, I., Sheiko, S.S., and Rubinstein, M. (2005) Diblock Copolymer Micelles in a Dilute Solution. Macromolecules, 38, 5330–5351.

24. Alexander, S. (1977) Adsorption of chain molecules with a polar head a scaling description. J. Phys. France, 28, 983–987.

25. de Gennes, P.G. (1980) Macromolecules, 13, 1069–1075.

26. Annianson, E.A.G. and Wall, S.N. (1974) On the Kinetics of Step-Wise Micelle Association. J. Phys. Chem., 78, 1024–1030.

27. Hadgiivanova, R., Diamant, H., and Andelman, D. (2011) Kinetics of Surfactant Micellization: A Free Energy Approach. J. Phys. Chem. B, 115, 7268–7280.

28. Mysona, J.A., McCormick, A.V., and Morse, D.C. (2019) Mechanism of Micelle Birth and Death. Phys. Rev. Lett., 123, 038003.

29. Ohta, T. and Kawasaki, K. (1986) Equilibrium Morphology of Block Copolymer Melts. Macromolecules, 19, 2621–2632.

30. Kawasaki, K., Ohta, T., and Kohrogui, M. (1988) Equilibrium Morphology of Block Copolymer Melts. 2. Macromolecules, 21, 2972–2980.

31. Uneyama, T. and Doi, M. (2005) Calculation of the Micellar Structure of Polymer Surfactant on the Basis of the Density Functional Theory. Macromolecules, 38, 5817–5825.

