## Supplementary Files for "Triblock copolymer micelle model of spherical paraspeckles"

### Supplementary Files: Triblock copolymer micelle model of spherical paraspeckles

E-mail:

#### S1 Free energy due to conformational entropy of B blocks

The free energy due to the conformational entropy of blocks in the core was derived by Semenov in the limit of strong segregation.[1] This theory treats the core as a spherical melt brush. We here use the same approach to analyze the free energy due to the conformational entropy of B blocks in the core of a paraspeckle. In a paraspeckle, B blocks of copolymers that have A blocks in the shell form loops, whereas B blocks of copolymers that have A blocks in the core are straight in the core. The free energy due to the conformational entropy has the form

$$\begin{aligned}\frac{F_{\rm c}}{k_{\rm B}T} &= \frac{3}{2b^2}\int_0^{r_{\rm c}} dr_0 \int_0^{r_0} dr\, E_{\rm s}(r,r_0)g_{\rm s}(r_0) \\ &+\frac{3}{b^2}\int_0^{r_{\rm c}} dr_0 \int_0^{r_0} dr E_{\rm l}(r,r_0)g_{\rm l}(r_0), \qquad \text{(S1)}\end{aligned}$$

where $E_{\rm s}(r,r_0) = |dr_{\rm s}(n,r_0)/dn|$ and $E_{\rm l}(r,r_0) = |dr_{\rm l}(n,r_0)/dn|$ are the local stretching of the straight and loop blocks. $r$ is the distance from the interface between the core and the shell. $g_{\rm s}(r_0)$ is the distribution functions of the free end of the straight blocks and $g_{\rm l}(r_0)$ is the distribution function of the middle segment of the loop blocks. With eq. (S1), we assume that the most probable path of a loop is symmetric with respect to the segment in the middle to the B block.

Eq. (S1) should be minimized with respect to $E_{\rm s}(r,r_0)$ and $E_{\rm l}(r,r_0)$ with the constraints due to the fact that all blocks have the same the number of segments

$$\int_0^{r_0} \frac{dr}{E_{\rm s}(r,r_0)} = N_{\rm A} + N_{\rm B} \qquad \text{(S2)}$$

$$2\int_0^{r_0} \frac{dr}{E_{\rm l}(r,r_0)} = N_{\rm B} \qquad \text{(S3)}$$

and the incompressibility condition

$$\int_r^{r_{\rm c}} dr_0 \frac{g_{\rm s}(r_0)}{E_{\rm s}(r,r_0)} + 2\int_r^{r_{\rm c}} dr_0 \frac{g_{\rm l}(r_0)}{E_{\rm l}(r,r_0)} = \frac{S_d(r_{\rm c}-r)}{b^d}. \qquad \text{(S4)}$$

The function $S_d(r)$ depends on the dimensionality $d$ ($d = 3$ for spherical micelles, $d = 2$ for cylindrical micelles, and $d = 1$ for lamellar) and has the form

$$S_1(r) = 2 \quad \text{(S5)}$$

$$S_2(r) = 2\pi r \quad \text{(S6)}$$

$$S_3(r) = 4\pi r^2. \quad \text{(S7)}$$

We take into account these constraints by using the Lagrange multipliers

$$\begin{aligned}
\frac{F_\mathrm{c}}{k_\mathrm{B}T} &= \frac{3}{2b^2}\int_0^{r_\mathrm{c}} dr_0 \int_0^{r_0} dr\, E_\mathrm{s}(r,r_0) g_\mathrm{s}(r_0) \\
&+\frac{3}{b^2}\int_0^{r_\mathrm{c}} dr_0 \int_0^{r_0} dr E_\mathrm{l}(r,r_0) g_\mathrm{l}(r_0) \\
&+\int_0^{r_\mathrm{c}} dr_0 \mu_\mathrm{s}(r_0) \int_0^{r_0} dr \frac{1}{E_\mathrm{s}(r,r_0)} \\
&+2\int_0^{r_\mathrm{c}} dr_0 \mu_\mathrm{l}(r_0) \int_0^{r_0} dr \frac{1}{E_\mathrm{l}(r,r_0)} \\
&+\int_0^{r_\mathrm{c}} dr \Pi_0(r) \int_r^{r_\mathrm{c}} dr_0 \left[\frac{g_\mathrm{s}(r_0)}{E_\mathrm{s}(r,r_0)} + \frac{2g_\mathrm{l}(r_0)}{E_\mathrm{l}(r,r_0)}\right].
\end{aligned} \quad \text{(S8)}$$

Applying the variational principle to eq. (S8) with respect to $E_\mathrm{s}(r,r_0)$ and $E_\mathrm{l}(r,r_0)$ leads to the form

$$E_\mathrm{s}^2(r,r_0) = \frac{2b^2}{3}\left(\Pi_0(r) + \frac{\mu_\mathrm{s}(r_0)}{g_\mathrm{s}(r_0)}\right) \quad \text{(S9)}$$

$$E_\mathrm{l}^2(r,r_0) = \frac{2b^2}{3}\left(\Pi_0(r) + \frac{\mu_\mathrm{l}(r_0)}{g_\mathrm{l}(r_0)}\right). \quad \text{(S10)}$$

The free ends of A blocks of unlooped chains are not stretched, $E_\mathrm{s}(r_0,r_0) = 0$. The turning segments of B blocks of loops are not stretched too, $E_\mathrm{l}(r_0,r_0) = 0$. These boundary conditions lead to the form

$$E_\mathrm{s}(r,r_0) = (\varphi_\mathrm{s}(r_0) - \varphi_\mathrm{s}(r))^{1/2} \quad \text{(S11)}$$

$$E_{\rm l}(r,r_0) \;=\; (\varphi_{\rm l}(r_0)-\varphi_{\rm l}(r))^{1/2}. \tag{S12}$$

Substituting eqs. (S11) and (S12) into eqs. (S2) and (S3) leads to the forms

$$\int_0^{r_0} dr \frac{1}{(\varphi_{\rm s}(r_0)-\varphi_{\rm s}(r))^{1/2}} \;=\; N_{\rm A}+N_{\rm B} \tag{S13}$$

$$2\int_0^{r_0} dr \frac{1}{(\varphi_{\rm l}(r_0)-\varphi_{\rm l}(r))^{1/2}} \;=\; N_{\rm B}. \tag{S14}$$

The solutions of eqs. (S13) and (S14) have the forms

$$E_{\rm s}(r,r_0) \;=\; \frac{\pi}{2N_{\rm B}}(r_0^2-r^2)^{1/2} \tag{S15}$$

$$E_{\rm l}(r,r_0) \;=\; \frac{\pi}{N_{\rm B}}(r_0^2-r^2)^{1/2}, \tag{S16}$$

where it is useful to refer to ref.[2] for the derivation. Substituting eqs. (S15) and (S16) into eq. (S4) leads to the form

$$\frac{2N_{\rm B}}{\pi}\int_r^{r_{\rm c}} dr_0 \frac{g_{\rm s}(r_0)+g_{\rm l}(r_0)}{\sqrt{r_0^2-r^2}} = \frac{S_d(r_{\rm c}-r)}{b^d}. \tag{S17}$$

For the case of spherical paraspeckle ($d=3$), the solution of eq. (S17) has the form

$$(N_{\rm A}+N_{\rm B})g_{\rm s}(r_0)+N_{\rm B}g_{\rm l}(r_0) = N_{\rm B}g(r_0), \tag{S18}$$

where $g(r_0)$ has the form

$$g(r_0) = \frac{8\pi r_{\rm c} r_0}{N_{\rm B} b^3}\left(\tanh^{-1}\sqrt{1-\frac{r_0^2}{r_{\rm c}^2}} - \sqrt{1-\frac{r_0^2}{r_{\rm c}^2}}\right). \tag{S19}$$

It is useful to refer to ref.[1] for the derivation of eq. (S19). The fraction of B blocks that form loops is $\alpha$,

$$\int_0^{r_0} g_{\mathrm{l}}(r_0) = \alpha. \tag{S20}$$

A simple summation scheme

$$g_{\mathrm{s}}(r_0) = (1-\alpha)\frac{N_{\mathrm{B}}}{N_{\mathrm{A}}+N_{\mathrm{B}}}g(r_0) \tag{S21}$$

$$g_{\mathrm{l}}(r_0) = \alpha g(r_0) \tag{S22}$$

satisfies both eqs. (S18) and (S20). Substituting eqs. (S21) and (S22) into eq. (S1) leads to the free energy due to the conformational entropy of B blocks,

$$\frac{F_{\mathrm{c}}}{k_{\mathrm{B}}T} = \frac{3}{2}\lambda_{\mathrm{s}} n \frac{r_{\mathrm{c}}^5}{N_{\mathrm{B}}^2 b^2}4\alpha + \frac{3}{2}\lambda_{\mathrm{s}} n \frac{r_{\mathrm{c}}^5}{(N_{\mathrm{A}}+N_{\mathrm{B}})^2 b^2}(1-\alpha), \tag{S23}$$

where $\lambda_{\mathrm{s}}$ $(=\pi^3/30)$ is a constant. The factor 4 in the first term of eq. (S23) reflects the fact that a loop of B block is viewed as two chains, each composed of $N_{\mathrm{B}}/2$ segments.

One criticism of this derivation may be that we used different unknown functions, $\varphi_{\mathrm{s}}(r)$ and $\varphi_{\mathrm{l}}(r)$, for the straight chains and loops in eqs. (S11) and (S12), although both of them are $2b^2\Pi_0(r)/3 +$ constant. Using different functions, $\varphi_{\mathrm{s}}(r)$ and $\varphi_{\mathrm{l}}(r)$, imply that different mean field pressure is applied to the segments of straight chains and loops. If we use the same function for the straight chains and loops, it is not possible to satisfy eqs. (S2) and (S3) whatever the choice of $g_{\mathrm{s}}(r_0)$ and $g_{\mathrm{l}}(r_0)$. The solutions, eqs. (S21) and (S22), ensure the constraints, eqs. (S2) - (S4), and thus are at least consistent. Another approach is to divide the core into two layers: one layer at the surface is composed of loops and unlooped chains and the other layer at the center is composed only of unlooped chains. However, we use eq. (S23) for the free energy due to the conformational entropy of B blocks.

### S2 Free energy of shell

#### S2.1 Mean field theory (derivation of eq. (4) in the main article

The free energy of the shell has the form

$$F_{\rm sh} = F_{\rm A} + F_{\rm C}, \tag{S24}$$

where $F_{\rm A}$ is the free energy of A domains and $F_{\rm C}$ is the free energy of C domains. A and C blocks are not mixed in the shell, but exist in different domains. We neglect the interactions between A and C blocks because the interactions are only between A and C blocks at the interface between domains.

The free energy of A domains has the form

$$\frac{F_{\rm A}}{k_{\rm B}T} = \int_{r_{\rm c}}^{r_{\rm A}} \frac{4\pi r^2 f_{\rm A} dr}{R_{\rm A}^3} \left[\frac{3}{2}\frac{R_{\rm A}^2(r)}{b^2 g_{\rm A}(r)} + v_{\rm A}\frac{g_{\rm A}^2(r)}{R_{\rm A}^3(r)}\right], \tag{S25}$$

where $R_{\rm A}(r)$ is the size of blobs at a distance $r$ from the center of the spherical paraspeckle and $g_{\rm A}(r)$ is the number of A segments in each blob. $r_{\rm c}$ is the radius of the core and $r_{\rm A}$ is the distance between the brush top of A domains and the center of the paraspeckle. $f_{\rm A}$ is the fraction of interfacial area occupied by A domains. $v_{\rm A}$ is the excluded volume that accounts for the repulsive interactions between A segments. $4\pi r^2 f_{\rm A}(r) dr/R_{\rm A}^3$ is the number of blobs in the spherical shell of thickness $dr$ at a distance $r$ from the center of the paraspeckle. The inside of the square bracket is the free energy of each blob and has the form of the free energy of the Flory theory of swollen chains. We here used mean field approximation to derive eq. (S25).

The free energy of C domains has a similar form

$$\frac{F_{\rm C}}{k_{\rm B}T} = \int_{r_{\rm c}}^{r_{\rm C}} \frac{4\pi r^2 (1-f_{\rm A}) dr}{R_{\rm C}^3} \left[\frac{3}{2}\frac{R_{\rm C}^2(r)}{b^2 g_{\rm C}(r)} + v_{\rm C}\frac{g_{\rm C}^2(r)}{R_{\rm C}^3(r)}\right], \tag{S26}$$

where $R_\mathrm{C}(r)$ is the size of blobs at a distance $r$ from the center and $g_\mathrm{C}(r)$ is the number of segments in each blob. $r_\mathrm{C}$ is the distance between the brush top of C domains and the center of the paraspeckle. $v_\mathrm{C}$ is the excluded volume that accounts for the repulsive interactions between C segments in the shell.

The Flory theory predicts that the relationship between the size $R_\mathrm{A}$ and the number $g_\mathrm{A}$ of segments has the form

$$R_\mathrm{A}(r) = b\left(\frac{v_\mathrm{A}}{b^3}\right)^{1/5} g_\mathrm{A}^{3/5}(r). \tag{S27}$$

The size $R_\mathrm{C}(r)$ and the number $g_\mathrm{C}(r)$ of segments has a similar relationship

$$R_\mathrm{C}(r) = b\left(\frac{v_\mathrm{C}}{b^3}\right)^{1/5} g_\mathrm{C}^{3/5}(r). \tag{S28}$$

The theory of polymer brush predicts that the size $R_\mathrm{A}(r)$ of blobs in A domains is determined by the average area occupied by one A block

$$4\pi r^2 f_\mathrm{A} = R_\mathrm{A}^2(r)\alpha n, \tag{S29}$$

where $\alpha$ is the fraction of A blocks in the shell. The size $R_\mathrm{C}(r)$ of blobs in C domains is determined by a similar manner

$$4\pi r^2(1 - f_\mathrm{A}) = R_\mathrm{C}^2(r)n. \tag{S30}$$

Eqs. (S27) - (S30) leads to the sizes, $R_\mathrm{A}$ and $R_\mathrm{C}$, and the number of segments, $g_\mathrm{A}(r)$ and $g_\mathrm{C}(r)$, once the fraction $f_\mathrm{A}$ is given.

The fraction $f_\mathrm{A}$ of area occupied by A domains is derived by the balance of the osmotic pressure between A domains and C domains,

$$v_{\rm A}\frac{g_{\rm A}^2(r)}{R_{\rm A}^6(r)} = v_{\rm C}\frac{g_{\rm C}^2(r)}{R_{\rm C}^6(r)}. \tag{S31}$$

Eqs. (S27) - (S31) leads to the form

$$f_{\rm A} = \frac{\alpha v_{\rm A}^{1/4}}{\alpha v_{\rm A}^{1/4} + v_{\rm C}^{1/4}} \tag{S32}$$

and it is not a function of the distance $r$.

The distances, $r_{\rm A}$ and $r_{\rm C}$, are derived by the conservation of the number of segments

$$\int_{r_{\rm c}}^{r_{\rm A}} \frac{4\pi r^2 f_{\rm A} dr}{R_{\rm A}^3(r)} g_{\rm A}(r) = N_{\rm A}\alpha n \tag{S33}$$

$$\int_{r_{\rm c}}^{r_{\rm C}} \frac{4\pi r^2 (1-f_{\rm A}) dr}{R_{\rm C}^3(r)} g_{\rm C}(r) = N_{\rm C} n. \tag{S34}$$

Eqs. (S33) and (S34) lead to the form

$$r_{\rm A} = r_{\rm c}\left(1 + \frac{5}{3}\frac{h_{\rm A}}{r_{\rm c}}\right)^{3/5} \tag{S35}$$

$$r_{\rm C} = r_{\rm c}\left(1 + \frac{5}{3}\frac{h_{\rm C}}{r_{\rm c}}\right)^{3/5}, \tag{S36}$$

where $h_{\rm A}$ and $h_{\rm C}$ have the form

$$h_{\rm A} = N_{\rm A} b\left(\frac{n\alpha}{4\pi f_{\rm A} r_{\rm c}^2}\frac{v_{\rm A}}{b}\right)^{1/3} \tag{S37}$$

$$h_{\rm C} = N_{\rm C} b\left(\frac{n}{4\pi(1-f_{\rm A}) r_{\rm c}^2}\frac{v_{\rm C}}{b}\right)^{1/3}. \tag{S38}$$

Eqs. (S37) and (S38) have the forms of the height of planer brushes (note that $\sigma_{\rm A} = n\alpha/(4\pi f_{\rm A} r_{\rm c}^2)$ and $\sigma_{\rm C} = n/(4\pi(1-f_{\rm A})r_{\rm c}^2)$ are the grafting density of A and C blocks, respectively). $h_{\rm A}$ and $h_{\rm C}$ are indeed the height of A and C domains in the limit of $r_{\rm c} \to \infty$, see eqs. (S35) and (S36).

By using eqs. (S27) - (S38), eqs. (S25) and (S26) are rewritten in the form

$$\frac{F_{\mathrm{A}}}{k_{\mathrm{B}}T} = \frac{15}{2}\alpha n \frac{h_{\mathrm{A}}}{bN_{\mathrm{A}}}\frac{r_{\mathrm{c}}}{b}\left[\left(1+\frac{5}{3}\frac{h_{\mathrm{A}}}{r_{\mathrm{c}}}\right)^{1/5}-1\right] \tag{S39}$$

$$\frac{F_{\mathrm{C}}}{k_{\mathrm{B}}T} = \frac{15}{2}n \frac{h_{\mathrm{C}}}{bN_{\mathrm{C}}}\frac{r_{\mathrm{c}}}{b}\left[\left(1+\frac{5}{3}\frac{h_{\mathrm{C}}}{r_{\mathrm{c}}}\right)^{1/5}-1\right]. \tag{S40}$$

In the limit of $r_{\mathrm{c}}$, eqs. (S39) and (S40) have approximate forms

$$\frac{F_{\mathrm{A}}}{k_{\mathrm{B}}T} = \frac{5}{2}\alpha n \frac{h_{\mathrm{A}}^2}{b^2 N_{\mathrm{A}}} \tag{S41}$$

$$\frac{F_{\mathrm{C}}}{k_{\mathrm{B}}T} = \frac{5}{2}n \frac{h_{\mathrm{C}}^2}{b^2 N_{\mathrm{C}}}, \tag{S42}$$

which are the forms of the free energy of planar brushes.

#### S2.2 Scaling theory

In a more precise treatment, the free energy contributions of A and C domains are the thermal energy $k_{\mathrm{B}}T$ per blob,

$$\frac{F_{\mathrm{A}}}{k_{\mathrm{B}}T} = C_{\mathrm{s}}\int_{r_{\mathrm{c}}}^{r_{\mathrm{A}}} \frac{4\pi r^2 f_{\mathrm{A}} dr}{R_{\mathrm{A}}^3(r)} \tag{S43}$$

$$\frac{F_{\mathrm{C}}}{k_{\mathrm{B}}T} = C_{\mathrm{s}}\int_{r_{\mathrm{c}}}^{r_{\mathrm{C}}} \frac{4\pi r^2 (1-f_{\mathrm{A}}) dr}{R_{\mathrm{C}}^3(r)}, \tag{S44}$$

where $C_{\mathrm{s}}$ is a constant of order unity and $C_{\mathrm{s}} \simeq 1.38$ was derived by curvefitting the results on polystylene-polyisoprene with the theory of diblock copolymer micelles.[3] By using eqs. (S27) - (S38), eqs. (S43) and (S44) are rewritten in the forms

$$\frac{F_{\mathrm{A}}}{k_{\mathrm{B}}T} = \frac{3}{5}\frac{(n\alpha)^{3/2}}{(4\pi f_{\mathrm{A}})^{1/2}} C_{\mathrm{s}} \log\left(1+\frac{5}{3}\frac{h_{\mathrm{A}}}{r_{\mathrm{c}}}\right) \tag{S45}$$

$$\frac{F_{\mathrm{C}}}{k_{\mathrm{B}}T} = \frac{3}{5}\frac{n^{3/2}}{(4\pi (1-f_{\mathrm{A}}))^{1/2}} C_{\mathrm{s}} \log\left(1+\frac{5}{3}\frac{h_{\mathrm{C}}}{r_{\mathrm{c}}}\right). \tag{S46}$$

The fraction $f_{\mathrm{A}}$ of the surface of the core occupied by A blocks is derived by the balance of the osmotic pressure

$$\frac{k_{\mathrm{B}}T}{R_{\mathrm{A}}^{3}(r)} = \frac{k_{\mathrm{B}}T}{R_{\mathrm{C}}^{3}(r)}, \tag{S47}$$

between the domains of A blocks and the domains of C blocks, where we used the fact that the pressure is the thermal energy per the volume of a blob. This leads to the form

$$f_{\mathrm{A}} = \frac{\alpha}{1+\alpha}. \tag{S48}$$

The heights, $h_{\mathrm{A}}$ and $h_{\mathrm{C}}$, in eq. (S46) have the same form as eqs. (S37) and (S38), respectively, but eq. (S48) should be used for $f_{\mathrm{A}}$.

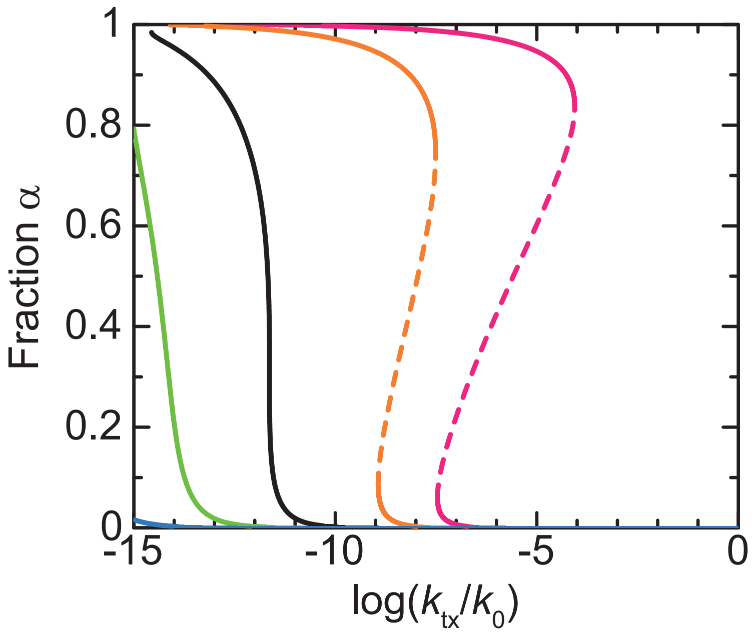


Figure S1: The fraction $\alpha$ of A blocks in the shell of a paraspeckle, as predicted by using the scaling theory, is shown as a function of the (natural) logarithm of the transcription rate $k_{\text{tx}}$ for $N_{\text{A}} = 1.0$ (cyan), 3.0 (light green), 5.08455 (black), 8.0 (orange), and 10.0 (magenta). We used $N_{\text{B}} = 40.0$, $N_{\text{C}} = 20.0$, $\chi_{\text{B}} = 0.5$, $\chi_{\text{AB}} = 1.0$, $\lambda_{\text{s}} = \pi^3/30$, and $C_{\text{s}} = 1.5$ for the calculations.

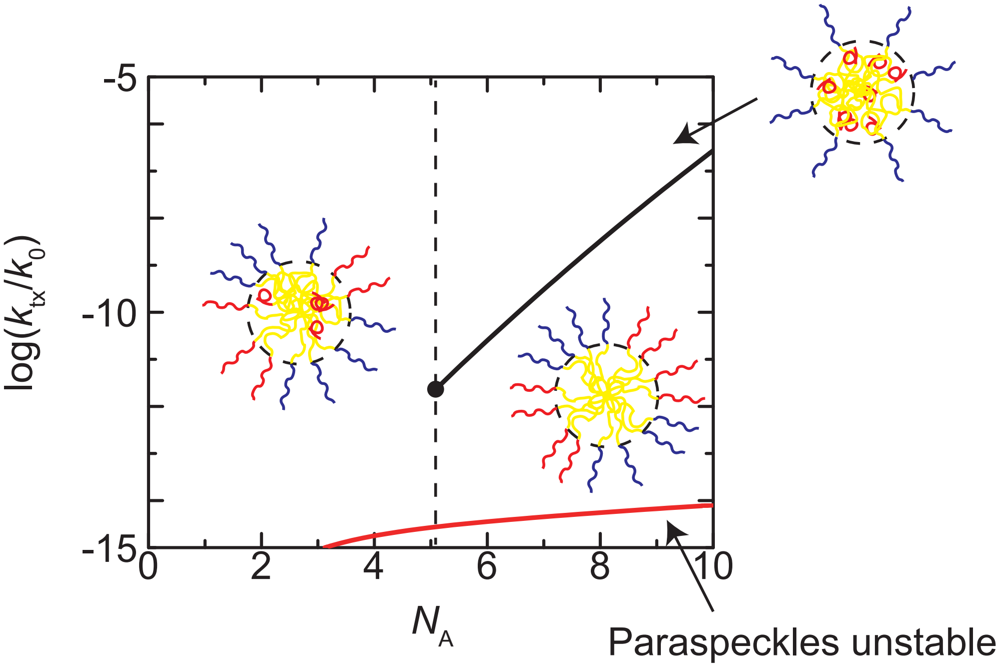


Figure S2: The phase diagram of paraspeckles, as predicted by using the scaling theory, is shown for the (natural) logarithm of production rate $k_{\rm tx}$ of copolymers and the number $N_{\rm A}$ of segments of A blocks. Paraspeckles are not stable in the region delineated by the red line. We used $N_{\rm B} = 40.0$, $N_{\rm C} = 20.0$, $\chi_{\rm B} = 0.5$, $\chi_{\rm AB} = 1.0$, $\lambda_{\rm s} = \pi^3/30$, and $C_{\rm s} = 1.5$ for the calculations.

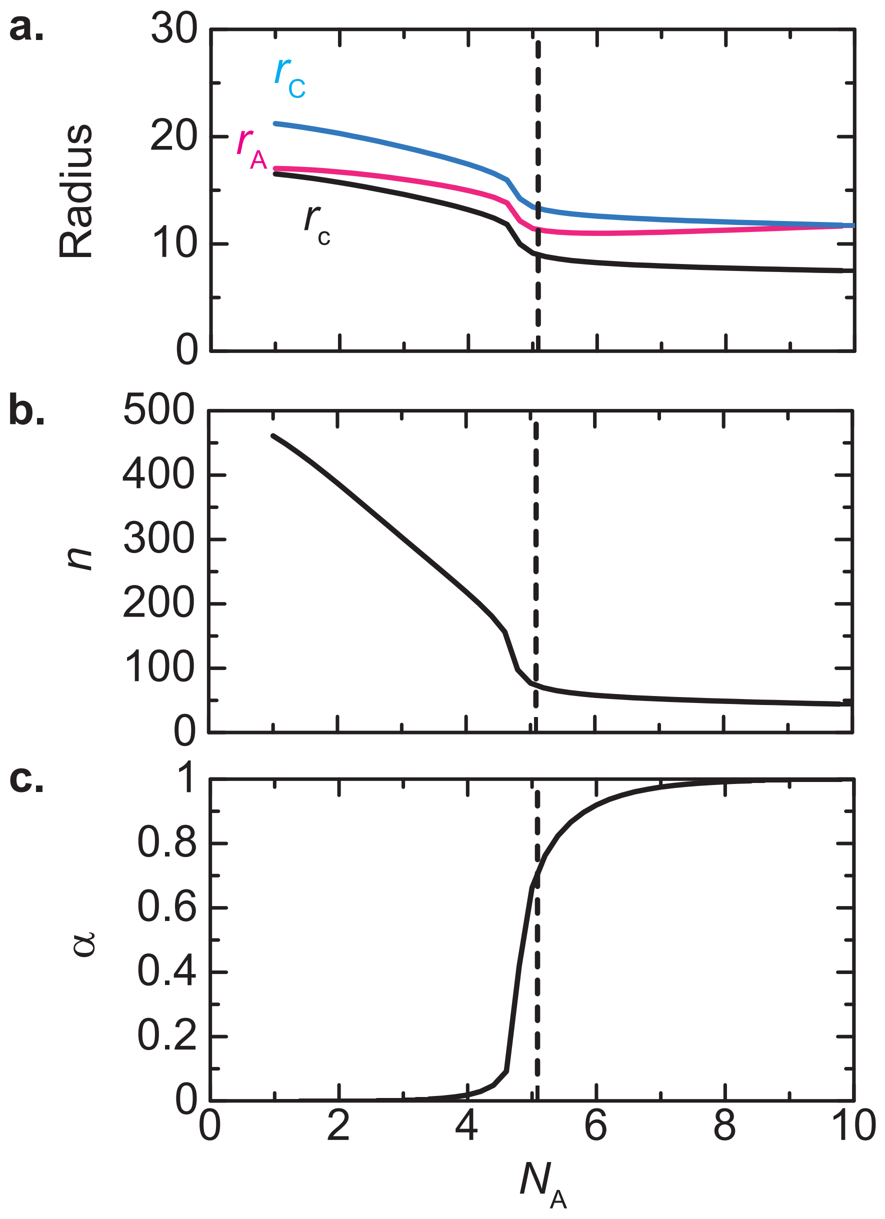


Figure S3: The radius (**a**), the number $n$ of copolymers (**b**), and the fraction $\alpha$ of A blocks in the shell of paraspeckles, as predicted by using the scaling theory, is shown as a function of the number $N_{\mathrm{A}}$ of segments of A blocks. In **a**, we showed the radius of the core of a paraspeckle (black), the distance between the top of a A domain and the center of the paraspeckle (magenta), and the distance between the top of a C domain and the center of the paraspeckle (cyan). The (natural) logarithm $\log(k_{\mathrm{tx}}/k_0)$ of the production rate of copolymers is fixed to $-12.0$. We used $N_{\mathrm{B}} = 40.0$, $N_{\mathrm{C}} = 20.0$, $\chi_{\mathrm{B}} = 0.5$, $\chi_{\mathrm{AB}} = 1.0$, $\lambda_{\mathrm{s}} = \pi^3/30$, and $C_{\mathrm{s}} = 1.5$ for the calculations.

### References

(1) Semenov,A.N. (1985) Contribution to the theory of microphase layering in block-copolymer melts. *Sov. Phys. JETP*, **61**, 733–742.

(2) Netz,R.R. and Schick,M. (1998) Polymer Brushes: From Self-Consistent Field Theory to Classical Theory. *Macromolecules*, **31**, 5105–5122.

(3) Zhulina,E.B., Adam,M., LaRue,I., Sheiko,S.S., and Rubinstein,M. (2005) Diblock Copolymer Micelles in a Dilute Solution. *Macromolecules*, **38**, 5330–5351.
